# Wearable Dual-Modality Plethysmography for Arterial Modulation and Blood Pressure Dip

**DOI:** 10.64898/2026.04.17.719282

**Authors:** Seobin Jung, Seamus D. Thomson

## Abstract

Continuous, non-invasive cardiovascular monitoring is limited by the superficial sensing depth of Photoplethysmography (PPG), which is susceptible to peripheral artifacts. This study evaluates a wearable dual-modality prototype integrating dry-electrode Impedance Plethysmography (IPG) and PPG within a smartwatch form factor. Results from a pilot study (N=2) demonstrate that IPG signals exhibit a temporal lead over PPG across ventral and dorsal sites, supporting its greater penetration depth. During brachial artery modulation, IPG showed superior sensitivity to arterial recovery on the ventral forearm. Furthermore, 60-minute napping sessions revealed that while PPG remained morphologically stable, IPG signals underwent significant evolution, capturing distinct pulsewave archetypes. These findings suggest that wearable IPG provides a high-fidelity window into deep systemic hemodynamics typically reserved for clinical instrumentation.

## I. Introduction

The landscape of cardiovascular health is undergoing a paradigm shift, moving from intermittent clinical assessments to continuous, non-invasive wearable technologies that provide longitudinal, real-world data [1]. In a consumer market increasingly driven by artificial intelligence and health-focused features, there is a growing demand for higher-fidelity sensor inputs that can move beyond simple heart rate tracking to capture more complex systematic hemodynamics [1], [2]. Currently, the industry standard for consumer wearables is Photoplethysmography (PPG) which tracks heart rate and oxygen saturation by measuring light absorption in the peripheral microvasculature [3], [4]. However, PPG is inherently limited by its reliance on superficial skin perfusion, making it highly susceptible to motion artifacts, temperature-induced vasoconstriction, skin tone, and contact pressure [5], [6].

Unlike optical sensors, Impedance Plethysmography (IPG) measures changes in electrical impedance across tissue, providing a more direct assessment of volumetric blood flow [7], [8]. More recently, IPG has gained traction in wearable device research as a robust alternative to optical sensors for cuffless blood pressure estimation and long-term hemodynamic monitoring [9]–[12]. However, the fundamental sensing principles regarding its penetration depth and specific volumetric origins - particularly when applied to the distal forearm - remain poorly understood. Our previous work in temperature-mediated vasoconstriction suggests that IPG is capable of sensing deeper blood flow than PPG [13]. In our 2026 study,we found that the IPG signal had no statistically-significant changes due to peripheral vasoconstriction compared to PPG when measuring on the ventral forearm using medical-grade gel electrodes. While compelling, those findings are limited by a form factor unsuitable for consumer use: the use of wet conductive gels and ventral placement are not conducive to long-term, real-world wear.

To bridge the gap between a lab-based proof-of-concept and viable wearable technology appropriate for consumer products, this study investigates the feasibility of a dual-modality wearable designed within the form-factor constraints of a commercial smartwatch (the Google Pixel Watch 2). Unlike previous iterations, this device utilizes dry stainless-steel electrodes integrated into the backplate for dorsal wrist sensing, representing a substantial step forward toward longitudinal consumer applications.

Moving beyond hardware feasibility, this study leverages the prototype to provide further mechanistic evidence of the sensing depth advantages inherent to IPG over PPG. By analyzing lead-lag timing between the two modalities – a methodology analogous to multi-wavelength PPG [14] - we propose an arterial modulation model to evaluate the temporal dynamics of pulse collapse and recovery. We hypothesize that a sensor with a greater depth penetration depth will detect changes in brachial artery inflow more rapidly than a sensor restricted to superficial microvasculature. Using a “water hose” analogy - where the main trunk responds to an inflow cutoff before its smaller branches - we aim to demonstrate that IPG offers a more direct window into deep arterial activity than the peripheral perfusion captured by PPG.

Finally, we investigate the clinical utility of this enhanced sensing depth for monitoring nocturnal blood pressure (BP) dipping. While ambulatory blood pressure monitoring (ABPM) remains the gold standard for capturing the characteristic 10-20% nocturnal BP drop, its reliance on intermittent cuff inflation is inherently intrusive and often disrupts the sleep it seeks to characterize [15]–[17]. We evaluate the capacity of our wearable-derived IPG signals to non-invasively capture the physiological shifts associated with these BP transitions during rest. Daytime napping may be used as a surrogate for the BP changes observed in nocturnal monitoring, wherein substantial BP reductions can occur within the first hour of sleep onset [18], [19]. Furthermore, siestas have been shown to produce mean BP levels and dipping magnitudes comparable to nighttime sleep [20], [21]. By comparing simultaneous IPG, PPG, and ABPM measurements during these types of sessions, we aim to demonstrate that this prototype can observe complex hemodynamic processes.

## II. Methodology

### A. Prototype Design

The experimental platform utilized a modified commercial smartwatch form factor (Google Pixel Watch 2) designed for dorsal wrist placement (Fig. 1A). To enable dual-modality sensing, the device was equipped with a custom-engineered backplate (Fig. 1B):

- IPG Module: A linear four-electrode array was integrated using stainless steel 316L. The outer current-injecting electrodes (I_1_, I_2_) each measured 100 mm^2^ (22 mm length, 4.55 mm width), while the inner voltage-sensing electrodes (V_1_, V_2_) measured 80 mm^2^ (22 mm length, 3.64 mm width). A 1.5 mm gap separated the injecting/sensing pairs, with an 18 mm center-to-center distance between the sensing electrodes. Electrodes featured a 0.25 mm protrusion to maintain consistent skin contact during wear.
- PPG Module: A reflective infrared PPG module (an off-the-shelf OPB733TR module from TT Electronics, England) was positioned at the geometric center of the electrode array to capture superficial microvascular perfusion. This sensor module was tested against the Biopac infrared PPG module SS4LA (Biopac Systems, CA, USA) for its optical equivalency to capture PPG pulsatiles.
- Structural Integration and Signal Routing: The housing was constructed from anodized aluminium and the backplate was made using Accura AMX Rigid Black (3D Systems, Inc., SC, USA). Wires (24 gauge, flexible silicone) were anchored via a VeroBlackPlus RGD875 (Stratasys Ltd., MN, USA) dome and interfaced with a custom Acrylonitrile Butadiene Styrene (ABS) plastic connection box to wire-out analog signals while ensuring signal integrity. The connection box contained a custom printed circuit board populated with a passive network for the PPG module such as a resistor for LED driver (1.5 kOhm) and a resistor for photodiode biasing (10 kOhm), and routing for both IPG and PPG to the data acquisition system.

**Fig. 1.**
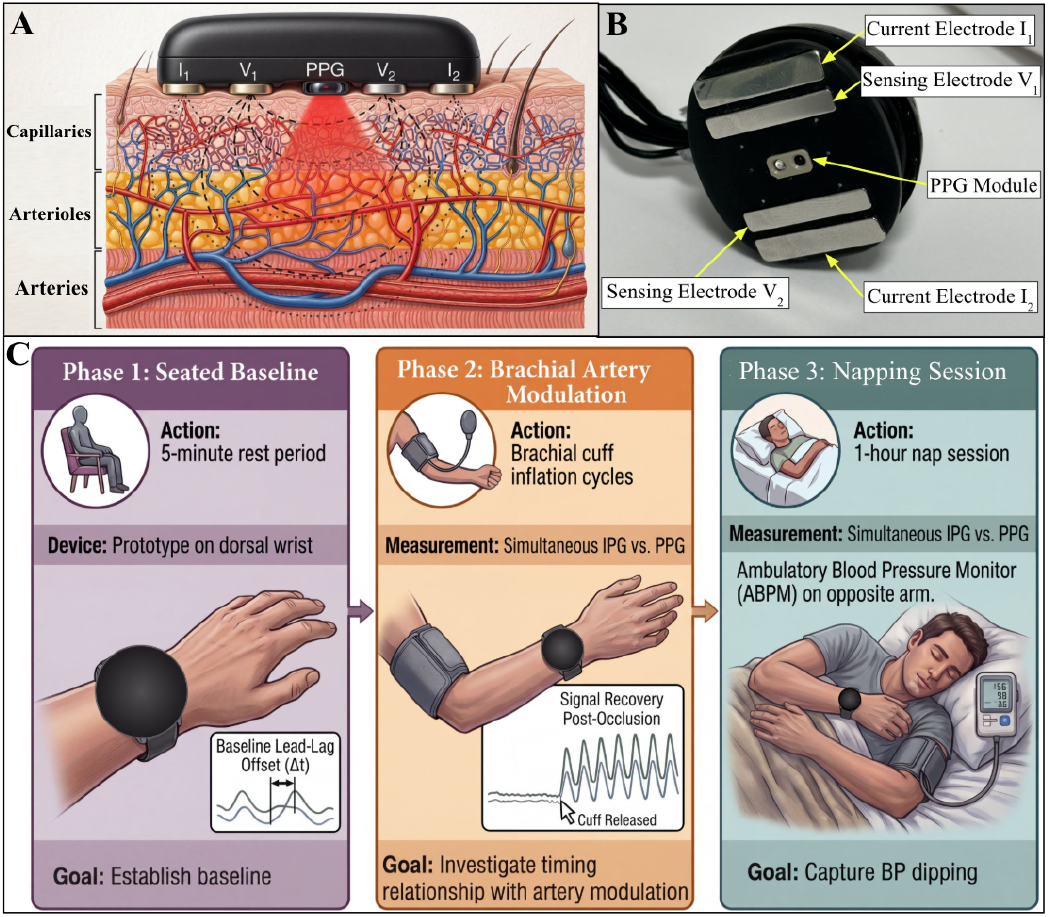
Cross sectional illustration (not to scale) of the dual-modality (IPG and PPG) wearable prototype positioned over the skin (A). Underside photograph of the prototype showing the positioning of IPG electrodes and PPG module (B). Illustration of the three-phase methodology evaluating the prototype feasibility (C).

### B. Data Acquisition and Signal Preprocessing

Synchronous IPG and PPG data were acquired using a Biopac MP36R system. IPG and PPG signals were acquired using the SS31L cardiac output module (100 kHz excitation frequency, excitation current = 400 µArms, continuously applied) and SS4LA module respectively. For prototype data collections, the SS4LA module optical component was substituted with the module described in 2.1 Prototype Design. For the non-prototype-based PPG data collection, the Biopac PPG module was used as it was. For ventral forearm collection, Electrocardiogram (ECG) signals were also recorded using gel electrodes configured for lead I (left shoulder, right shoulder, and right leg ankle) using Biopac SS2LB.

- Sampling Rates: Data were sampled at 2 kHz for Phases 1 and 2. A data rate of 250 Hz was selected for Phase 3 due to the data volume associated with this longer collection.
- Signal Filtering: Both modalities underwent an 8th-order zero-phase Butterworth bandpass filter (0.5 - 10 Hz).
- Z-Channel Derivation: To mitigate saturation in the primary impedance (Z) channel, the first derivative (dZ/dt) was integrated to reconstruct the volumetric Z signal. Baseline wandering was corrected for Phase 3 using a moving average subtraction (window = 250 samples); this was to improve fiducial point extraction during this longitudinal collection. Baseline correction was not applied to Phase 1 and 2 as the arterial modulation intervention was believed to reflect baseline shifts.

### C. Experimental Design

Two healthy adult participants were recruited for this feasibility investigation. The study protocol was divided into three distinct phases to evaluate sensor performance across varying physiological states (Fig. 1C):

- Phase 1 Seated Baseline: Participants remained seated at a table and for a five-minute period to establish baseline pulsatile morphology archetypes and characterize the lead-lag offset between IPG and PPG. They were instructed to relax, trying not to move or talk during the data collection. 60 seconds of IPG and PPG signal were acquired and used for this baseline measurement. For ground truth comparison, the same protocol was also performed using IPG with 3M 2760-5 gel electrodes on the ventral forearm and the Biopac PPG module - following a similar positioning to our earlier work [13].
- Phase 2 Brachial Artery Modulation: To evaluate the sensitivity of the volumetric IPG and PPG signals to changes in vascular resistance, arterial flow was modulated using a standard automated brachial arm cuff BP monitor (Omron, Japan). This occurred immediately after Phase 1. Participants were seated with their arms resting on a table. The arm cuff was positioned on their left arm (the same side as the prototype). Brachial artery flow was modulated by inflating the BP cuff and release, a method shown to impact this blood flow [22]. Multiple cycles of cuff inflation (occlusion) and release (recovery) were performed to evaluate the collapse and recovery timing of the IPG and PPG signals. The timing of data collection occurred as follows: 60 seconds of baseline (i.e. Phase 1), before the automated cuff inflation started. After the cuff was released/deflated, 30 seconds would pass before the next cycle began. A total of three cycles occurred per participant per measurement location. This protocol was also performed using IPG with 3M gel electrodes on the ventral forearm, along with the Biopac PPG module, to serve as a ground truth for vascular kinetics.
- Phase 3 Napping Session: Participants were transitioned to a supine position in a controlled environment and instructed to remain relaxed for at least 60 minutes. ABPM was performed using an ABPM7100 (WelchAllyn, USA) on the contralateral arm to the prototype at regular 20-minute intervals to track systolic (SBP), diastolic (DBP), and mean arterial pressure (MAP) as ground truth for the nocturnal dip.

All experimental procedures were performed in accordance with the Declaration of Helsinki. The study protocol was approved by the Institutional Review Board of the WIRB-Copernicus Group (Study ID: 1391946). All participants provided written informed consent prior to the commencement of data collection, including specific consent for the publication of de-identified physiological waveforms.

### D. Data Analysis

#### Data were analyzed to focus on primary metrics for each phase using Python

- Phase 1 Temporal Dynamics: Fiducial points such as pulse onsets, systolic peaks, and maximum of second time derivatives (a-points) were identified using the pyPPG library [23]. The lead-lag offset (Δt shown in Fig. 1C) was calculated as the time difference from a PPG fiducial point to a corresponding IPG fiducial point. Pulse onsets were more erroneously identified than a-points; as a result, a-points were used to calculate the lead-lag offset.
- Phase 2 Volumetric Sensitivity: For the artery modulation data from Phase 2, for an ease of comparing timing relationships during pulse collapse and pulse recovery, IPG and PPG signals were first normalized using average amplitude values from respective baselines, and their root mean square (RMS) values were calculated in a rolling manner (window size = 3 sec).
- Phase 3 Morphological Heterogeneity: To ensure the integrity of the cardiac cycles to be used for morphological analysis, a multi-step beat cleaning procedure was implemented. Initial pulsewave segments were refined by recalculating onsets as the absolute minimum value between the consecutive u-points, which represents the points of maximum systolic slope identified using a first-derivative Savitzky–Golay filter with a window length of 9 samples and a polynomial order of 3. Following this segmentation, an outlier rejection step was applied based on beat duration; segments were rejected if their length fell outside the range of 50% to 150% of the median beat interval. Next, each validation beat underwent linear detrending (tilt correction) by subtracting a linear fit using the first and last points of the segment to remove low-frequency baseline wander. This systematic approach produced a refined set of high-quality beats for subsequent dimensionality reduction and clustering.

The cleaned pulsewave signals were processed using the scikit-learn library, with individual beats first resampled to a fixed length of 256 samples using cubic spline interpolation from the scipy.interpolate module. To ensure numerical stability during the decomposition, each beat was standardized using Z-score normalization, where the mean was subtracted and the result divided by the standard deviation plus a small number (10^-9^). The PCA implementation utilized sklearn.decomposition.PCA class to identify the principal components of the standardized beat vectors. Rather than using a fixed number of com-ponents, the dimensionality was reduced dynamically by calculating the cumulative explained variance ratio and selecting the minimum number of components required to reach a 95% threshold. This optimized feature set, representing the core morphological characteristics, was then used as the input for a k-means clustering algorithm and a silhouette scoring to identify distinct pulsewave archetypes.

## III. Results

### A. Baseline Morphology and Temporal Lead-Lag (Phase 1)

The prototype-based data acquisition from the dorsal wrist yielded clear pulsatiles for both IPG and PPG (Fig. 2A). Visual analysis revealed that IPG pulses exhibited a steeper systolic upstroke and a more pronounced dicrotic notch compared to the PPG signal. A persistent temporal offset was observed between the two modalities. The IPG pulse onset led the PPG onset, with mean time delta values from PPG a-point to IPG a-point (Participant 1, Participant 2): gel ventral forearm (-34.2 ms, -7.2 ms); dry electrode prototype on dorsal wrist (0.2 ms,-31.0 ms), and median values (Participant 1, Participant 2): gel ventral forearm (-71.0 ms, -13.5 ms); dry electrode prototype on the dorsal wrist (-4.5 ms, -16.3 ms) (Fig. 2B).

**Fig. 2.**
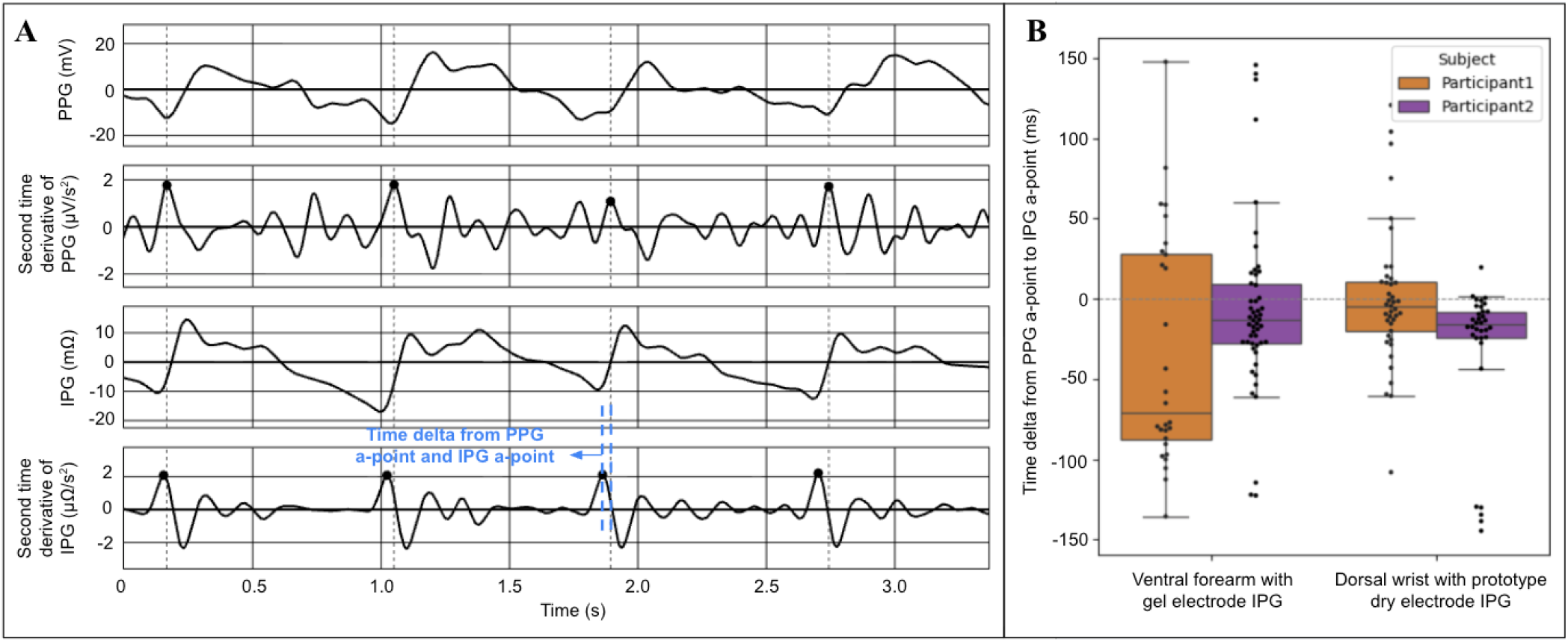
Simultaneously-acquired IPG and PPG from the wearable prototype showing time difference offset (A). Aggregated time differences from PPG a-point to IPG a-point for the two participants (B).

### B. Vascular Kinetics During Brachial Artery Modulation (Phase 2)

The “water hose” analogy was first evaluated using gel electrodes on the ventral forearm through cuff-induced occlusion and release. The Biopac PPG module was located in between the inner sensing gel electrode for IPG. Visual inspection of signal quality yielded poor signal quality for Participant 1 PPG; therefore, insights from this participant were limited. For Participant 2, both sensors tracked the cessation of pulsatility during full occlusion (Fig. 3A). First cuff inflation and deflation show that IPG and PPG pulses collapsed simultaneously (Fig. 3B) while IPG pulses recovered before PPG pulses recovered (Fig. 3C). Looking at the rolling RMS plot (Fig. 3D), it was observed that IPG and PPG signal collapse occurred at the same time for two repeats; the third repeat IPG signal collapsed after PPG. For signal recovery, IPG led PPG for all three repeats. Following the residual pressure from the cuff, IPG pulsatility returned to ∼50% baseline amplitude faster than PPG. The IPG signal reached a stable post-occlusion steady state within 3-4 cardiac cycles.

**Fig. 3.**
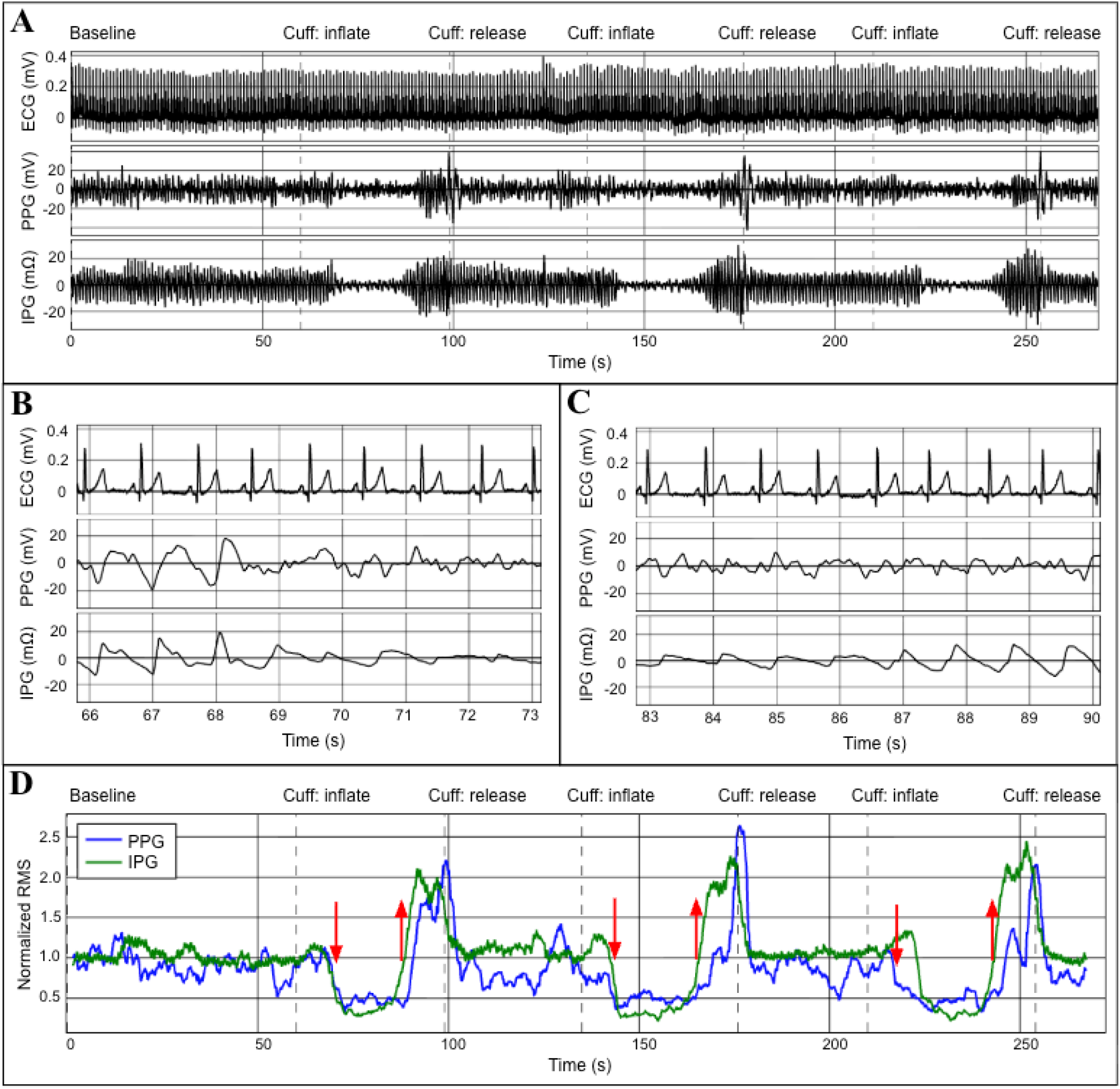
Full experimental trace of ECG, PPG, and IPG signals during three cycles of cuff-induced occlusion and release on the ventral forearm. Zoomed view of pulse collapse during cuff inflation, where both signals typically disappear in unison (B). Zoomed view of pulse recovery post-release, showing IPG pulsatility returning faster than PPG (C). Rolling normalized RMS plot illustrates that pulse recovery timing (upwards arrows) was led by IPG while pulse collapse timing (downwards arrows) was mixed (D).

For the dry electrode prototype positioned on the dorsal wrist, visual inspection of IPG and PPG signal quality revealed clear beats for both participants. The IPG and PPG signal collapse and recovery overlapped across all three repeats and for both participants; i.e. neither signal appeared or disappeared before the other.

### C. Napping Session (Phase 3)

Participants were supine throughout the entire napping session which successfully captured the characteristic nocturnal BP dip in both participants, as confirmed by the contralateral ABPM (Fig. 4). Participant 1 showed a maximum SBP reduction of 9 mmHg (7.6%) and a MAP reduction of 10 mmHg (10.3%) from baseline. Participant 2 exhibited an SBP reduction of 8 mmHg (7.5%) and a MAP reduction of 5 mmHg (5.9%). These hemodynamic shifts were accompanied by a simultaneous decrease in heart rate (HR), with maximum drops of 12 bpm (13.8%) and 3 bpm (5%) for participants 1 and 2, respectively. There was a correlation between HR and SBP for both participants (Fig. 4D).

**Fig. 4.**
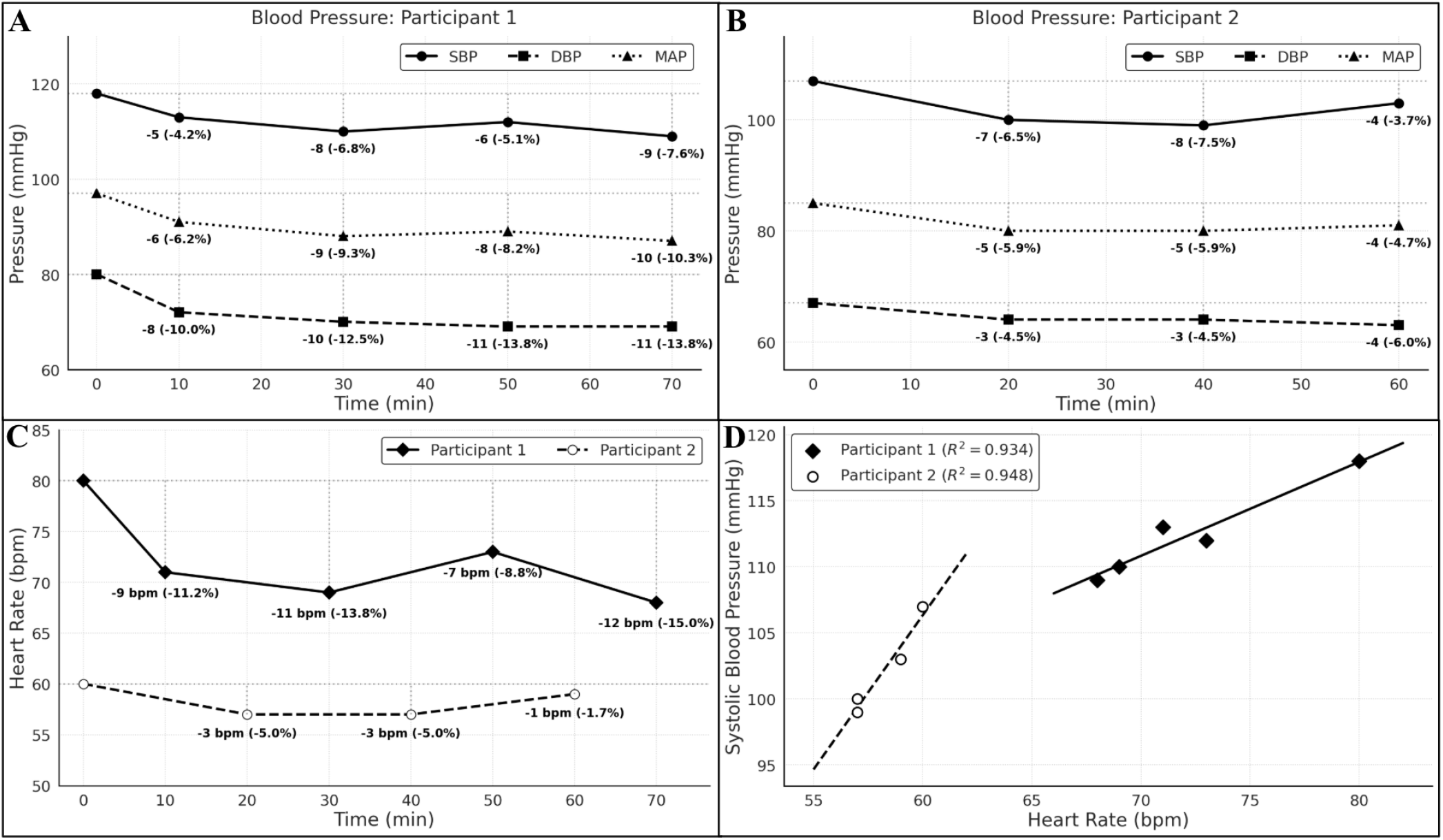
Blood pressure trends (SBP, DBP, and MAP) for Participant 1 and Participant 2 as measured by contralateral ABPM (A-B). Concurrent heart rate (HR) reductions observed during the 60-minute rest period for both participants (C). Linear correlation between HR and SBP (R^2^ *>* 0.93 for both participants) (D).

As the session progressed, a divergence in waveform morphology was observed between the two modalities best shown in Participant 1’s data (Fig. 5). At the 5-minute baseline (Fig. 5A), IPG and PPG pulsatiles appeared visually synchronous and morphologically similar. However, at subsequent snapshots (e.g. t = 20, 40, and 60 minutes) the IPG signal underwent profound evolution, characterized by the emergence of complex multi-peak features found in both the systolic and diastolic portions. In contrast, the PPG waveform remained largely consistent in its morphology.

**Fig. 5.**
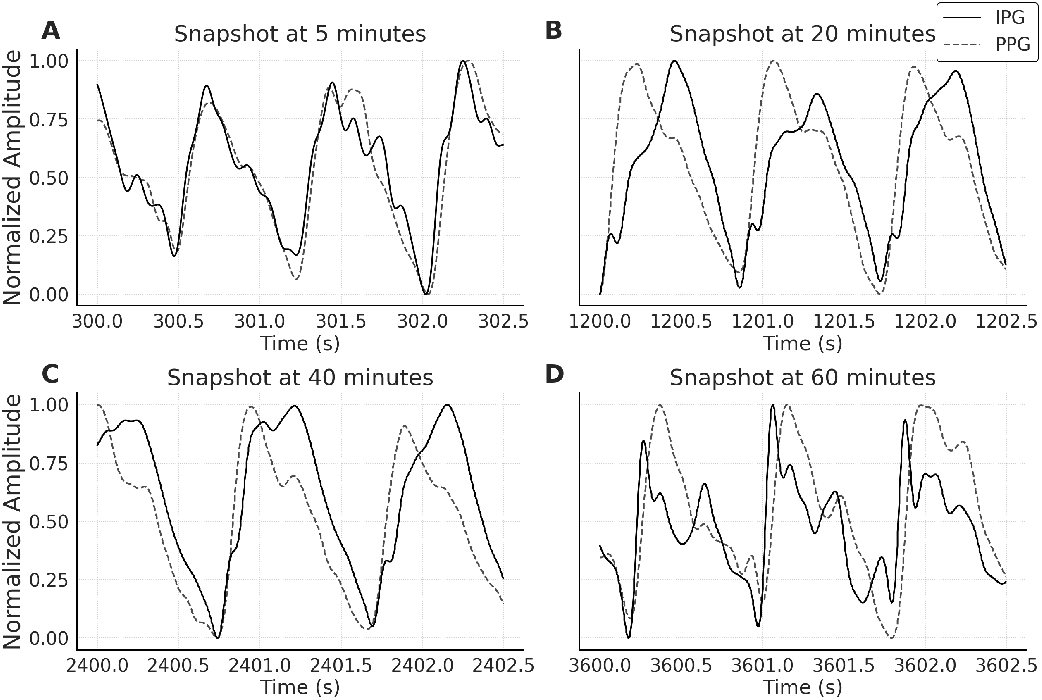
Snapshots of normalized heartbeat waveforms for Participant 1 taken at 5 minutes (A), 20 minutes (B), 40 minutes (C), and 60 minutes (D). While the PPG waveform remains morphologically stable throughout the session, the IPG signal undergoes evolution, transitioning from a simple pulse to a complex waveform with emergent multi-peak features in both the systolic and diastolic phases.

To quantify this heterogeneity, PCA and k-means clustering were applied to the heartbeat vectors of both modalities; revealing star contrast in signal dynamism (Fig. 6). For both participants, PPG data clustered into a single dominant mode, which maintained near-total prevalence (*>*95%) throughout the entire 60-minute collection. Conversely, IPG data revealed multiple distinct archetypes - three modes for Participant 1 and two modes for Participant 2. Temporal distribution of these modes (Fig. 6C-D) highlights the transitional nature of the IPG signal. For Participant 1, mode 3 (containing pronounced diastolic peaks) became the dominant archetype during the final 20% of the session. For Participant 2, mode 2 was more prevalent during the initial and final stages of the nap.

**Fig. 6.**
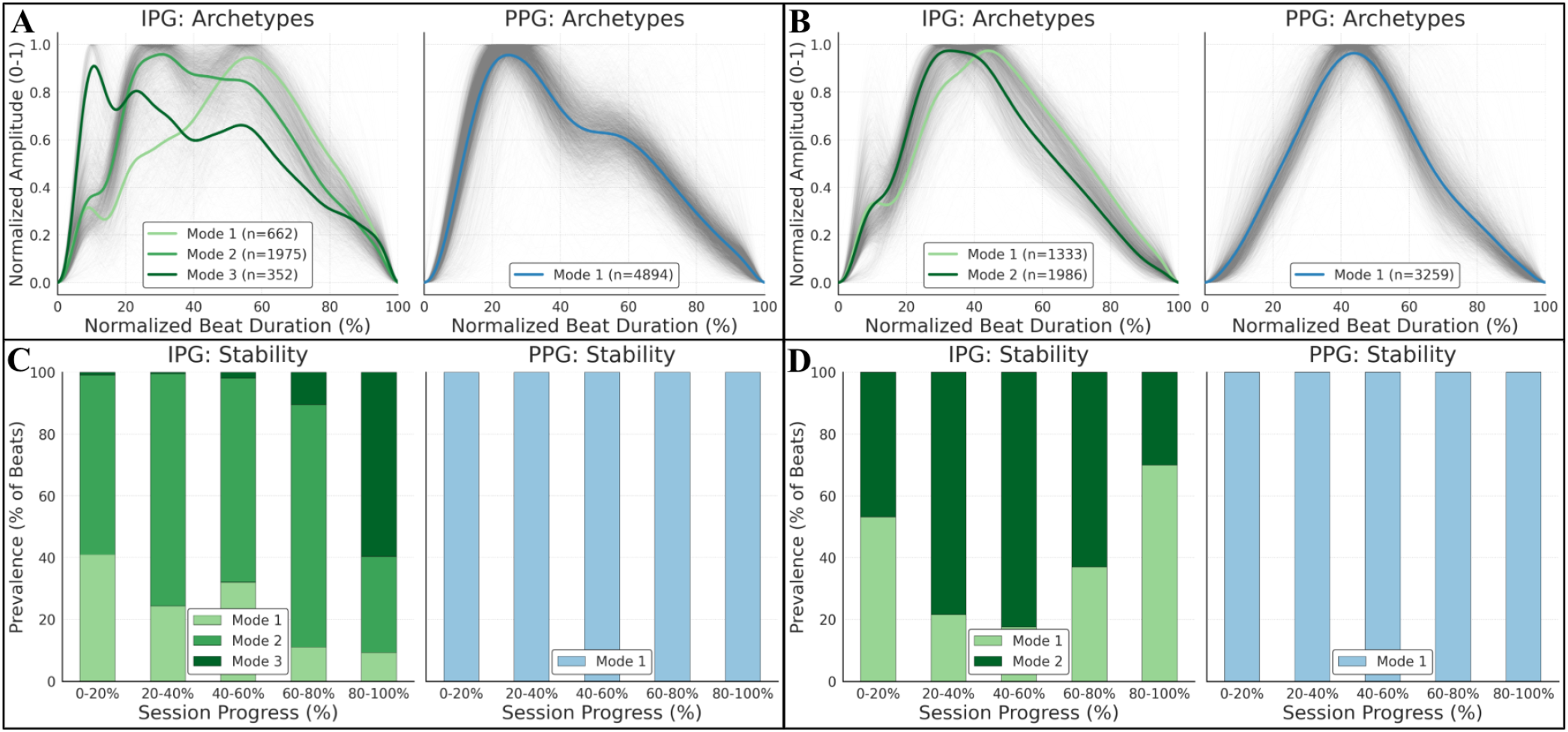
Identification of pulsewave archetypes for both participants. IPG data revealed multiple distinct morphological modes: three for Participant 1 (A) and two for Participant 2 (B), whereas PPG data remained restricted to a single dominant mode (*>* 95% prevalence). Temporal distribution of these modes throughout the napping session for Participant 1 (C) and Participant 2 (D).

## IV. Discussion

Our findings build upon earlier work which established that IPG provides a deeper sensing depth than PPG, making it inherently more resilient to peripheral vasoconstriction [13]. The transition from medical-grade gel electrodes to a dry stainless-steel interface within a commercial smartwatch form factor represents a critical step towards longitudinal health monitoring. We refined this dry electrode prototype through several iterations, involving experimentation with electrode areas, geometries, and alignments. For IPG sensing in a consumer form factor, the underside constraints of the Pixel Watch informed the prototype design, making a linear electrode array with the PPG module centered. While PPG remains the industry standard, its susceptibility to superficial artifacts often compromises signal quality in wrist-worn applications [5]. In contrast, the pulsatile IPG signals observed across multiple measurement sites in this pilot study demonstrate that dry-electrode bioimpedance can meet the signal-to-noise requirements of a wearable device without necessitating conductive gels.

The Phase 1 analysis evaluated the hypothesis that IPG signals would exhibit a consistent temporal lead over PPG. This negative lead-lag offset, persistent across both the ventral forearm gel electrode measurements and the dorsal wrist dry electrode prototype measurements, suggests that the IPG modality captures the arrival of the pressure wave in the deeper, larger-diameter arteries before the resulting perfusion reaches the superficial microvasculature. This hemodynamic latency could serve as a proxy for sensing depth, supporting the notion that IPG provides a more direct assessment of systemic arterial activity than the peripheral perfusion measured by PPG. McMurray et al. in their 2023 paper have also observed this latency between IPG and PPG using a linear gel electrode prototype, suggesting that this offset may be due to a forward compression of cells, fibers, and anatomical structures proceeding the cardiac pulse [26]. While our data corroborates with these latency patterns, further systematic research is required to fully disentangle the specific biomechanical and electro-physical variables at play within the complex, heterogeneous environment of the distal forearm.

For Phase 2, we hypothesized that the greater sensing depth of IPG would allow for the earlier detection of arterial inflow collapse and recovery during brachial artery modulation compared to superficial microvascular measurements of PPG. Our pilot results (N=2), however, were mixed and highly dependent on measurement location. On the ventral forearm, we observed that IPG pulsatility recovered before PPG in one participant, while the timing of the pulse collapse remained inconclusive; data for the second participant was unusable due to poor PPG signal quality. In contrast, IPG and PPG signals from the dorsal wrist moved in unison during both occlusion and recovery for both participants. These findings suggest that while brachial artery modulation induces systemic vascular changes, the localized sensitivity of IPG to deep arterial pulsatility is governed by regional anatomy. On the ventral forearm, the closer proximity of the radial and ulnar arteries to the skin surface likely facilitates a more direct current path through these major blood vessels, enhancing the sensor’s ability to resolve independent arterial kinetics. Conversely, the dorsal wrist is characterized by a more complex landscape of bone and connective tissue, which may attenuate the sensor’s sensitivity to these vessels, thereby explaining the lack of a temporal lead-lag during occlusion and recovery cycles at this site.

Evaluation of the napping session data (Phase 3) provides a critical proof-of-concept for the utility of wearable IPG in capturing BP dipping hemodynamics. Both participants exhibited a blood pressure drop relative to their baseline, but only Participant 1 displayed the characteristic 10% or greater reduction in SBP and MAP associated with a “dipper” profile [24]. Blood pressure is a multi-factorial product of cardiac output and systemic vascular resistance; while HR and SBP were correlated, other physiological variables likely contribute to this decline. A pivotal finding of this study is the observed divergence in waveform morphology as participants transitioned into deeper states of rest. While the PPG signal remained morphologically homogenous throughout the session, the IPG waveforms exhibited profound evolution, evidenced by the emergence of distinct archetypes for Participant 1 (Fig. 5). PCA-based clustering quantified this heterogeneity, revealing multiple IPG modes compared to a singular, stable PPG mode for both participants (Fig. 6). We interpret this disparity as further evidence of the differing sensing depths between the two modalities. As nocturnal dipping involves complex mechanisms such as vasodilation and sympathetic tone [25], the IPG signal likely captures the resulting changes in pulse wave reflections and arterial wall tension more effectively than PPG. Specifically, as the session progressed for Participant 1, the third mode (consisting of several peaks in the diastolic beat phase) became dominant in the last 20% of the collection. For Participant 2, their mode 2 was prevalent at the beginning and end of the collection, coinciding with ABPM readings where BP dropped at 20 and 40 minutes but began returning to baseline by the 60-minute mark. This temporal distribution may reflect the latencies associated with sleep onset and premature arousal. The ability to resolve these unique IPG archetypes during a 60-minute session is particularly noteworthy given the limitations of current ABPM tools, demonstrating that dry-electrode IPG may track these transitions non-invasively and continuously.

We acknowledge the inherent limitations of this feasibility study, specifically the small sample size (N=2) which precludes broad statistical generalization. The variability in mode prevalence between participants is a unique finding of this work, and likely stems from individual differences in autonomic regulation and peripheral vascular geometry. Further studies should validate these IPG archetypes against gold-standard instrumentation gold-standard instruments (e.g. continuous invasive BP monitoring or polysomnography) to definitively map specific morphological modes to sleep stages and hemodynamic states. Nevertheless, this study demonstrates that integrating dual-modality IPG-PPG sensing into a smartwatch consumer form factor provides a high-fidelity window into cardiovascular health, potentially moving complex hemodynamic monitoring from the clinical to the wrist.

## Acknowledgments

S.J. and S.T. contributed equally to the conceptualization, hardware design, experimental protocol for artery modulation, data analysis, and manuscript preparation. The authors would like to thank Keith Wong for his help in making the dual-modality plethysmography prototype. The illustrations of Fig. 1A and C were refined using Google Gemini.

